# Hypoxia Tolerance Declines with Age in the Absence of Methionine Sulfoxide Reductase (MSR) in *Drosophila melanogaster*

**DOI:** 10.1101/2021.06.02.446794

**Authors:** Nirthieca Suthakaran, Sanjana Chandran, Michael Iacobelli, David Binninger

## Abstract

Unlike the mammalian brain, *Drosophila melanogaster* can tolerate several hours of hypoxia without any tissue injury by entering a protective coma known as spreading depression. However, when oxygen is reintroduced, there is an increased production of reactive oxygen species (ROS) that causes oxidative damage. Methionine sulfoxide reductase (MSR) acts to restore functionality to oxidized methionine residues. In the present study, we have characterized in vivo effects of MSR deficiency on hypoxia tolerance throughout the lifespan of *Drosophila*. Flies subjected to sudden hypoxia that lacked MSR activity exhibited a longer recovery time and a reduced ability to survive hypoxic stress as they approached senescence. However, when hypoxia was induced slowly, MSR deficient flies recovered significantly quicker throughout their entire adult lifespan. In addition, the wildtype and MSR deficient flies had nearly 100% survival rates throughout their lifespan. Neuro-protective signaling mediated by decreased apoptotic pathway activation, as well as gene reprogramming and metabolic downregulation are possible reasons for why MSR deficient flies have faster recovery time and a higher survival rate upon slow induction of spreading depression. Our data are the first to suggest important roles of MSR and longevity pathways in hypoxia tolerance exhibited by *Drosophila*.

## 1. Introduction

Tolerance to a diminished level of oxygen (hypoxia) is a complex process that leads to a variety of responses by different organisms. The mammalian brain only tolerates a few minutes of severe hypoxia before it causes irreversible cell damage [1]. In contrast, *Drosophila melanogaster* has evolved a mechanism through which it can tolerate several hours of hypoxia without significant tissue injury. It does this by entering a protective coma known as spreading depression. However, when oxygen is reintroduced by reperfusion, there is an increased production of reactive oxygen species (ROS).

ROS are a group of highly reactive molecules that contain oxygen including radicals, molecules with unpaired valence electrons. Examples of ROS are superoxide (O_2_^−^), hydrogen peroxide (H_2_O_2_), and the hydroxyl radical (•OH) [2]. The accumulation of ROS is an unavoidable byproduct of cellular respiration, primarily within the mitochondrial electron transport chain. Superoxide is formed from the partial reduction of oxygen due to electron leakage at complex I and III of the electron transport chain [3]. ROS can also be produced through external sources such as UV light and radiation. Some ROS molecules lead to oxidative stress by oxidizing macromolecules and other cellular components, causing functionality loss [3]. For example, ROS oxidize the sulfur atom of methionine residues, oxidizing it to methionine sulfoxide (met-(o)) and often leading to the loss of protein function [4,5]. However, at low levels, certain ROS molecules, such as hydrogen peroxide, can also have a positive function since they regulate core metabolic pathways and play a role in signaling [3,6]. To prevent oxidative damage, cells can destroy the ROS molecules before they harm cellular components with the help of specific enzymes. Superoxide dismutase (SOD) is a well-studied example. SOD, which is present in the mitochondria and cytoplasm, converts the superoxide ion into hydrogen peroxide. Glutathione peroxidase (GPX) then reduces the hydrogen peroxide to water [3].

Oxidative damage can be reversed. Methionine sulfoxide reductase (MSR) represents a family of enzymes that have the primary role of reversing oxidative damage by reducing met-(o) back to its original form. The first evidence for MSR activity was described by Weissbach, Brot and colleagues when they observed that oxidation of methionine residues that inactivated ribosomal protein L12 in *Escherichia coli* [7] could be restored by MSRA [8]. There are two forms of MSR designated MSRA and MSRB, which are responsible for the stereospecific reduction of met-(o). MSRA reduces the S enantiomer (met-S-(o)) while MSRB reduces the R enantiomer (met-R-(o)). MSR also reverses oxidative damage by functioning as a scavenger of ROS to prevent damage to cellular components [9]. Mammals including humans have one *MSRA* gene and three distinct *MSRB* genes (*MSRB1*, *MSRB2*, *MSRB3*). *Drosophila* also has a single *MSRA* gene, but it only has one *MSRB* gene.

According to the free radical theory of aging, the accumulation of ROS and other free radicals plays a significant role in aging by causing oxidative damage [10]. Previous studies have detected increased ROS production in aged tissues, emphasizing the link between oxidative damage and aging [11]. In addition, met-(o) levels in proteins have been shown to increase with age in several aging models, such as replicative senescence and erythrocyte aging. Decreased levels of MSRA have been found in aged mouse tissues, and MSRA and MSRB were found to be downregulated during replicative senescence of human WI-38 fibroblasts [12]. Therefore, it is evident that there is a relationship between oxidative stress, the level of MSR activity, and aging. However, further studies are necessary to fully understand the role of MSR in aging and the onset of neurodegenerative diseases.

This study examines the effects of MSR in aging for *Drosophila melanogaster*. More specifically, it investigates the effect of MSR-deficiency on hypoxia tolerance throughout the lifespan of the adult *Drosophila*.

## 2. Materials and Methods

### 2.1 Fly Stocks

*Drosophila* stocks were maintained on standard cornmeal agar medium (Genesee Scientific, El Cajon, CA USA) at 25°C with a 12-hour light/dark cycle. The WT60 strain is homozygous wild-type for both the *MSRA* and *MSRB* loci (*MSRA*^+/+^*MSRB*^+/+^). The AB46 strain is fully deficient for any MSR activity due to homozygous null alleles for both the *MSRA* and *MSRB* genes (*MSRA*^Δ/Δ^MSR*B*^Δ/Δ^). The generation and characterization of these genetic lines has been described [13].

### 2.2 Hypoxia Treatment

Male flies were maintained with 50 animals per vial. All experiments were done between 12:00-5:00 PM to minimize changes in behavior affected by circadian rhythms. Chronic hypoxia was induced and maintained by continuous displacement of air in the hypoxia chamber (63.5 cm long × 33.7 cm wide × 43.8 cm high) with nitrogen gas (100%) at the calculated flow rate using Weymouth’s formula [14]. The rapid nitrogen flow rate was calculated as 2.0L/sec., the moderate flow rate was 1.4L/sec and the slow flow rate was 25mL/sec. An outlet in the chamber prevented any pressure changes within the chamber. A Pasco PASPort oxygen sensor (Pasco; Roseville, CA USA) was used to confirm that the concentration of oxygen inside the chamber was between 0-5% regardless of the nitrogen flow rate. The hypoxic conditions were maintained for the indicated time and nitrogen flow rate. Flies were then returned to standard atmosphere and monitored for recovery from the spreading depression coma.

### 2.3 Monitoring Fly Movement

Animal movement was monitored using a Drosophila Activity Monitor (DAM; TriKinetics Inc, Waltham, MA USA) which uses 32 glass tubes in an 8 × 4 array. One fly is placed in each tube which has small holes to allow free exchange of gasses. An infrared detector monitors the movement of the fly, recording each time the animal moves through the beam of the detector. Each trial began with an acclimation period of 10 minutes before data recording commenced. The onset of the spreading depression coma was measured as the time to the first cessation of movement detected by the DAM monitoring system. After the indicated time of hypoxic stress, flies were returned to normal atmosphere. The time to recovery was scored as the first movement of the animal detected by the DAM movement detection system. Flies that remained comatose after five hours of recovery were marked as failing to survive the hypoxia.

### 2.4 Statistical Analyses

Data were analyzed using Prism statistical software (GraphPad Software, San Diego, CA). The sample number (n) in each trial represents the number of flies, one in each glass tube, whose individual movements were monitored by the DAM movement detection system. Data points falling outside (outliers) the interquartile range (IQR) were not used in determining the average, standard error of the mean (SEM), sample number (n) and statistical significance.

## 3. Results

### 3.1. Characterization and Age-Grouping of Wild-Type and MSR-Deficient Strains

Complete loss of function (null) alleles of *MSRA* and *MSRB* were created by imprecise P-element transposon excision [13]. Briefly, the *MSRA* null allele is a 1.5kb deletion that starts 300bp upstream of the transcription start site and extends into Exon 2. The entire 5’ UTR and a portion of the open reading frame have been removed. The *MSRB* null allele is a 2.5kb deletion that starts 364bp upstream of the transcription start site and terminates 2163bp into the transcribed region. The loss of these genomic sequences removes the first three exons that includes a part of the open reading frame.

Strains that are homozygous wild-type for both *MSR* loci, homozygous for the *MSRA* deletion, homozygous for the *MSRB* deletion and MSR-deficient lines due to being homozygous for both deletions were used for this study. Experiments involving one hour of exposure to hypoxia were used for all four strains. However, no significant and reproducible differences between the wild-type strain and the strains that were homozygous for only one mutant locus were observed. Therefore, all the reported experiments only involve wild-type (WT60) and MSR-deficient (AB46) strains.

The absence of any functional *MSR* loci leads to nearly a 50% shorter lifespan compared to the wildtype [13]. Preliminary experiments showed that there were age-related differences in the response of the two strains to hypoxia. Animals were examined at a young age, middle age, and old age. Adjustments were made to the time period for each stage in their lifespan to account for the significantly shorter lifespan of the MSR-deficient line (Table 1).

**Table 1.** Categorization of Age Groups.

| Age Group | Wild-Type | MSR-Deficient |
| --- | --- | --- |
| Young | 20-25 Days Old | 5-10 Days Old |
| Middle Age | 40-45 Days Old | 30-35 Days Old |
| Old | 60-65 Days Old | 40-45 Days Old |

<sup>1</sup> *MSR*-deficient animals (AB46) have a markedly shorter lifespan. The ages selected to identify young, middle-age and old adult flies for each genotype are based on a previous study as described in the text.

### 3.2. Rapid Induction of the Spreading Depression Coma

A flow rate of 2.0L/sec nitrogen gas induced a spreading depression coma in both wild-type and MSR-deficient flies in less than one minute. Onset of the protective coma occurred too quickly to determine whether there was a difference between the two strains.

After one hour of acute hypoxia due to continuous air displacement with nitrogen, the animals were returned to normoxic (standard atmosphere) conditions and allowed to recover. Recovery was scored as the first movement detected by the infrared detector of the Drosophila Activity Monitor (see methods). There was an age-dependent effect in the recovery times for both wild-type and the MSR-deficient flies with an increase in recovery time as the animals aged (Figure 1). However, the MSR-deficient animals took significantly longer (t-test; p < 0.0001) compared to wild-type to recover at every age through their entire lifespan. This difference was most notable among the old animals where the wildtype took an average of 65.6 minutes to recover, but the MSR-deficient animals took 72% longer with an average of 112.8 minutes (p <0.0001).

**Figure 1.**
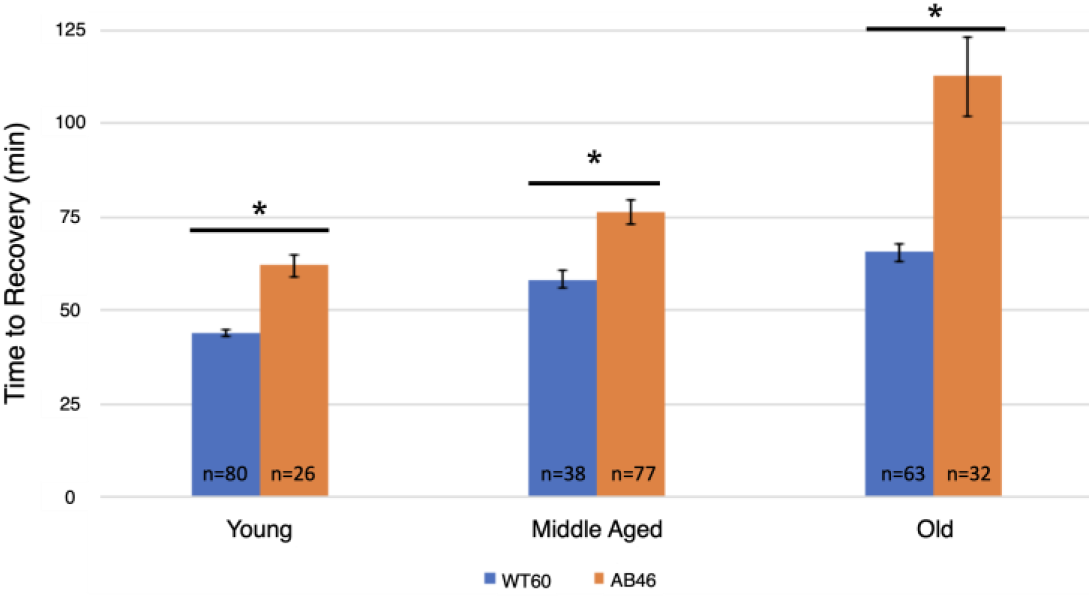
Recovery after One Hour Hypoxia Under Rapid Induction of Spreading Depression. Wild-type (WT60) and MSR-deficient (AB46) animals were exposed to one hour of hypoxia using a nitrogen flow rate of 2.0L/sec. After one hour of hypoxia, the animals were returned to normoxic conditions and allowed to recover from the spreading depression coma. Recovery was marked as the time to the first movement detected by the Drosophila Activity Monitor. The number of individual animals (n) used for each age-group and genotype is shown on the graph. Error bars are the SEM. Significance is indicated by an asterisk where p< 0.0001 using an unpaired t-test.

### 3.3. Moderate and Slow Induction of Hypoxic Coma

Two slower flow rates of nitrogen were used to extend the time to onset of the spreading depression coma to better determine whether there was an effect due to the absence of MSR activity. Onset of the coma was determined by the time required to no longer detect movement by the IR sensor of the Drosophila Activity Monitor. Overall, there was an age-dependent decrease in the time to induce the coma for both the moderate (Figure 2A) and slow (Figure 2B) flow rates of nitrogen gas. Interestingly, significant differences in the time for coma induction only occurred among middle age flies using the moderate flow rate and young flies using the slow flow rate. Wildtype middle age flies took an average of 4.1 minutes (n=45) whereas the MSR-deficient flies took 24% longer with an average of 5.1 minutes (n=120; p < 0.0001). Under the slow induction of spreading depression, wild-type flies took an average of 6.75 minutes (n=83) which was 28% longer than the MSR-deficient flies which took an average of 5.3 minutes (n=57; p < 0.0001).

**Figure 2.**
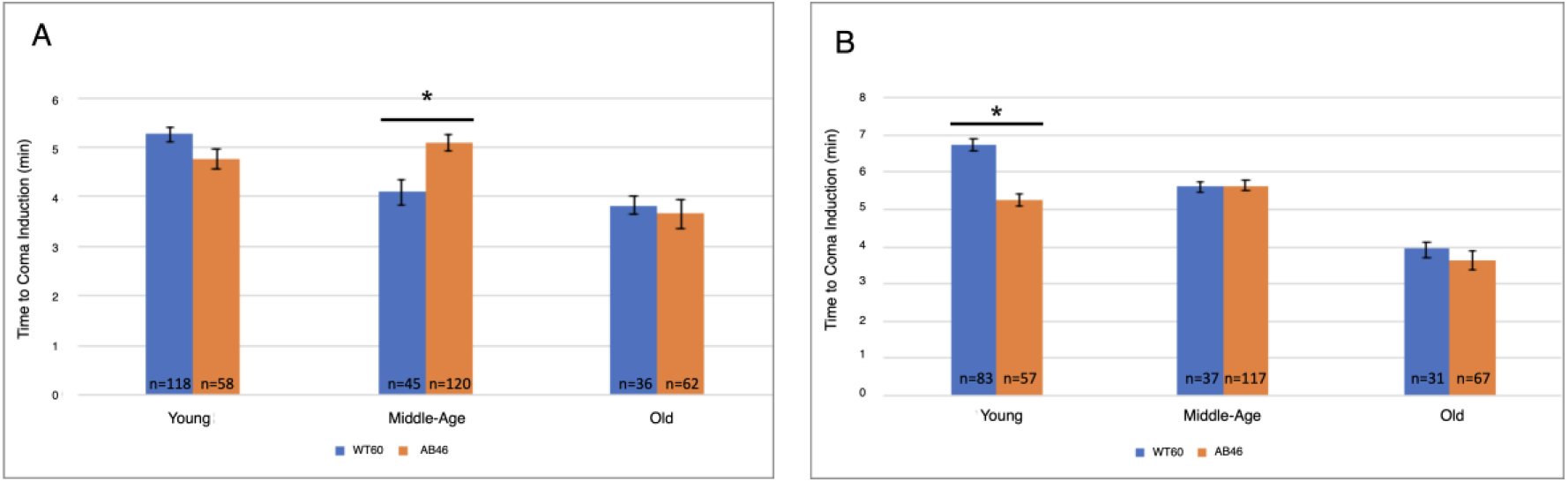
Moderate and Slow Induction of Spreading Depression. Wild-type (WT60) and MSR-deficient (AB46) animals were exposed to one hour of hypoxia. A nitrogen gas flowrate of 1.4L/sec was used for a moderate rate of induction (panel A) while a flow rate of 25mL/sec was used for a slow rate of induction (panel B). Onset of the coma was scored when the IR detector no longer recorded movement. The number of individual animals (n) used for each age-group and genotype is shown on the graph. Error bars are the SEM. Significance is indicated by an asterisk where p < 0.0001 using an unpaired t-test.

### 3.4. Recovery Following Moderate and Slow Induction of Hypoxia

The animals were returned to a normal atmosphere after one hour of hypoxic exposure to nitrogen gas at the indicated flow rate. Recovery was monitored as the first movement detected by the IR sensor as described above. There was an overall increase in recovery time for both wild-type and MSR-deficient animals as they aged. Unexpectedly, we found that the MSR-deficient animals recovered significantly faster than the wild-type at every age group for both the moderate (Figure 3A) and slow (Figure 3B) nitrogen flow rates. The difference was most pronounced in the old animals. When the coma was induced under a moderate nitrogen flow rate, old wild-type flies took an average of 127.2 minutes which was 73% longer than the average of 73.6 minutes required for recovery of the old MSR-deficient animals (p < 0.0001). The old wild-type animals recovered more quickly when the slow flow rate of nitrogen was used whereas recovery of the old MSR-deficient animals was nearly the same as those under a moderate nitrogen flow rate (73.6 minutes vs. 72.4 minutes). The wild-type flies still recovered more slowly with an average time of 108.0 minutes which was 49% longer than the average of 72.4 minutes required by the old MSR-deficient flies (p <0.0001).

**Figure 3.**
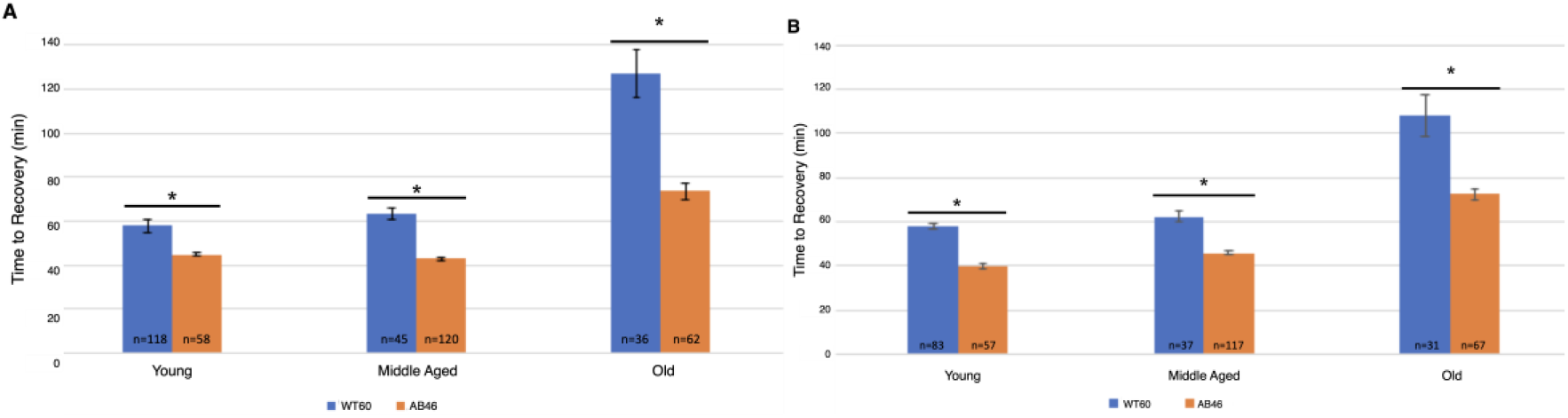
Recovery after One Hour Hypoxia Under Moderate or Slow Induction of Spreading Depression. Wild-type (WT60) and MSR-deficient (AB46) animals were exposed to one hour of hypoxia using a moderate nitrogen flow rate of 1.4L/sec (panel A) or a slow flow rate of 25mL/sec (panel B). After one hour of hypoxia, the animals were returned to standard atmosphere and allowed to recover, their activity was scored as the first movement detected by the Drosophila Activity Monitor. The number of individual animals (n) used for each age-group and genotype is shown on the graph. Error bars are the SEM. Significance is indicated by an asterisk where p< 0.0001 using an unpaired t-test.

### 3.5. Survival Following One Hour of Hypoxia

The flow rate used for inducing the spreading depression coma had a major effect on the survival of animals as they aged. Survival of the wild-type strain was 99-100% at all ages tested and under all three modes for inducing spreading depression (data not shown). In contrast, survival of the MSR-deficient strain (AB46) as the animals aged was strongly dependent on mode of coma induction. The young MSR-deficient animals had a 99-100% survival irrespective of the nitrogen flow rate, which was like the survival success of the wild-type strain (Figure 4). At middle-age, the MSR-deficient had a 93-99% survival when the moderate or slow flow rate was used but survival declined to 79% using the rapid nitrogen flow rate (Figure 4). The effect of the nitrogen flow rate on survival was most pronounced among the old MSR-deficient flies. Nearly all the animals (99%) survived with the slow nitrogen flow rate. Even at the moderate nitrogen flow rate, the survival rate was 93%. However, under the rapid flow rate, where the MSR-deficient animals took 72% longer than the wildtype to recover from the hypoxia (Figure 1), only 38% of the animals survived.

**Figure 4.**
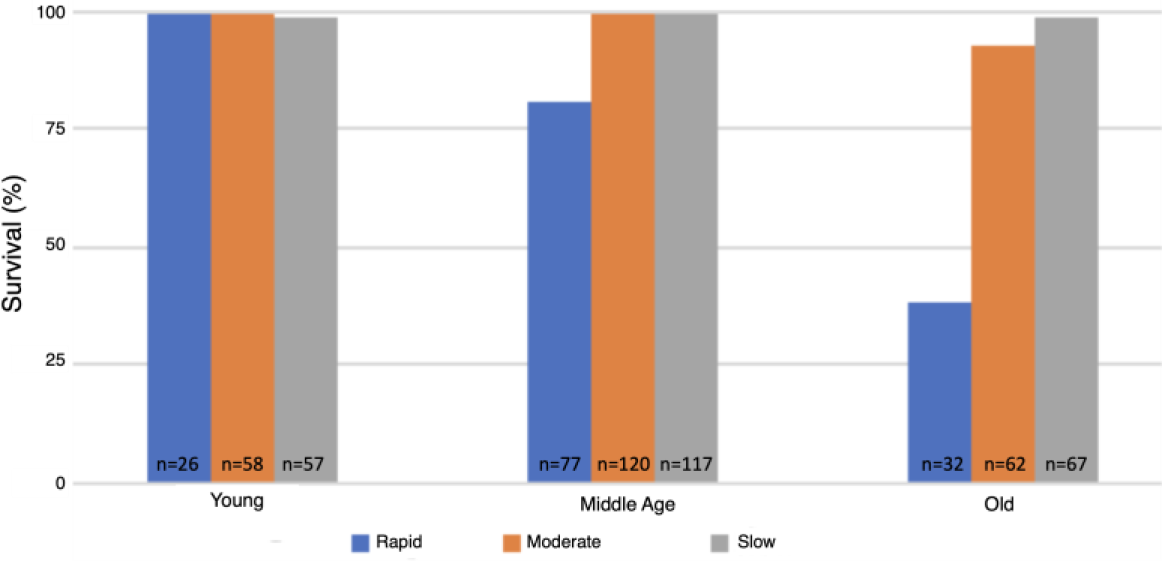
Survival Following Hypoxia of the MSR-Deficient Animals. The MSR-deficient (AB46) flies were subjected to 60 minutes hypoxia using the indicated flow rate of nitrogen. The flies were then returned to standard atmosphere and monitored for recovery from the coma as described in the legend of Figure 2. Animals that did not recover movement within 5 hours were scored as not surviving.

### 3.6. Effects of Prolonged Hypoxia on Recovery and Survival

All the previous experiments used one hour of hypoxic exposure. However, wild-type *Drosophila* can survive hours of chronic hypoxia [15,16]. Therefore, we examined the effect of three hours and six hours of hypoxia on the recovery and survival of the MSR-deficient flies. For these experiments, the slowest flow rate (25mL/sec) was used since the recovery and survival of the MSR-deficient flies was most robust.

Young MSR-deficient flies recovered significantly faster than the wild-type after three hours of hypoxia, using the slow flow rate of nitrogen (Figure 5A). Young wild-type flies took 98.2 minutes to recover which was 31% longer than the 75.5 minutes for recovery of the young MSR-deficient animals (p < 0.0001). After six hours of hypoxia (Figure 5B), the wild-type flies took twice as long to recover (197.0 minutes) compared to three hours of hypoxia. The MSR-deficient flies also took longer to recover (176.5 minutes) after the two additional hours of hypoxia (compare Figure 3 to Figure 5). While the MSR-deficient strains recovered faster than the wild-type after six hours of hypoxia, the difference was not significant (p = 0.08).

**Figure 5.**
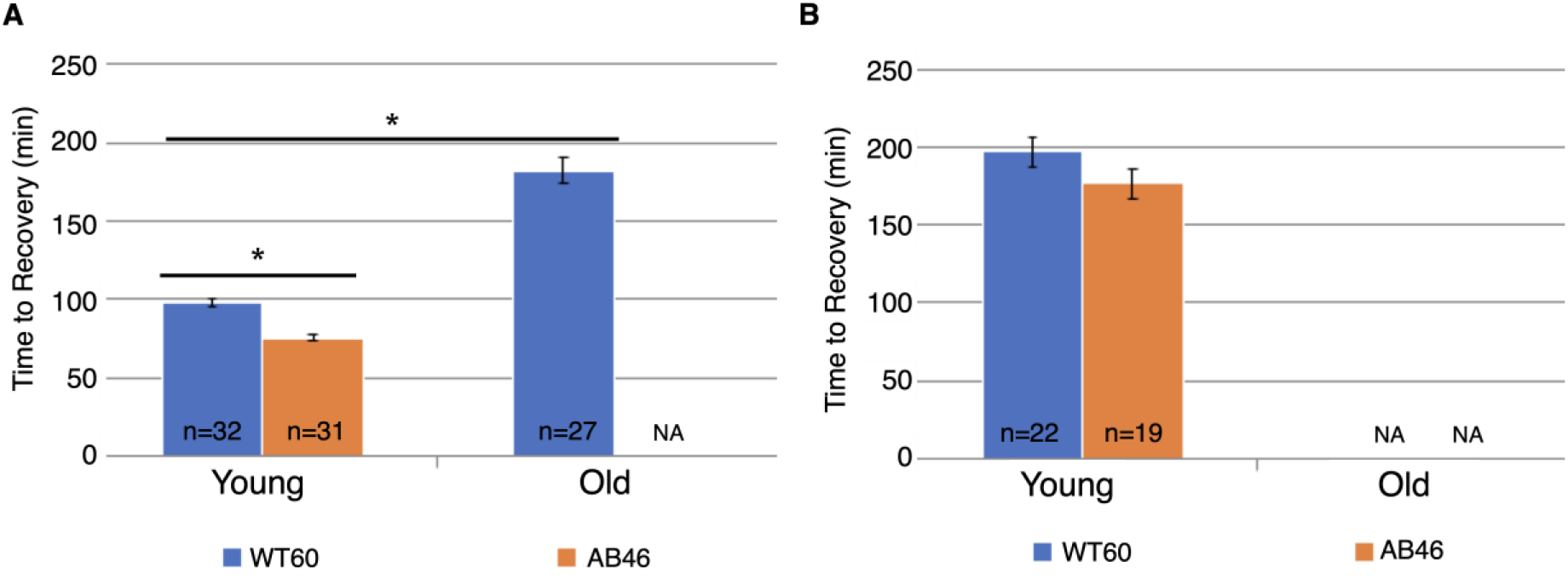
Recovery after Prolonged Hypoxia. Wild-type (WT60) and MSR-deficient (AB46) animals were exposed to three hours (panel A) or six hours (panel B) hypoxia using the slow nitrogen flow rate (25mL/sec). After the indicated time of hypoxia, the animals were returned to normal atmosphere and allowed to recover from the spreading depression coma. Recovery was scored as the first movement detected by the Drosophila Activity Monitor. The number of individual animals (n) used for each age-group and genotype is shown on the graph. NA indicates that none of the test animals survived the hypoxic treatment. Error bars are the SEM. Significance is indicated by an asterisk where p< 0.0001 using an unpaired t-test.

All the young test animals of both genotypes survived both 3 hours and 6 hours of hypoxia (Figure 6). However, there was a striking effect of age. All the old wildtype flies survived 3 hours of hypoxia whereas only 3% old MSR-deficient flies (1 of 32 animals) survived. In sharp contrast to the young animals, none of the old flies of either genotype survived six hours of hypoxia.

**Figure 6.**
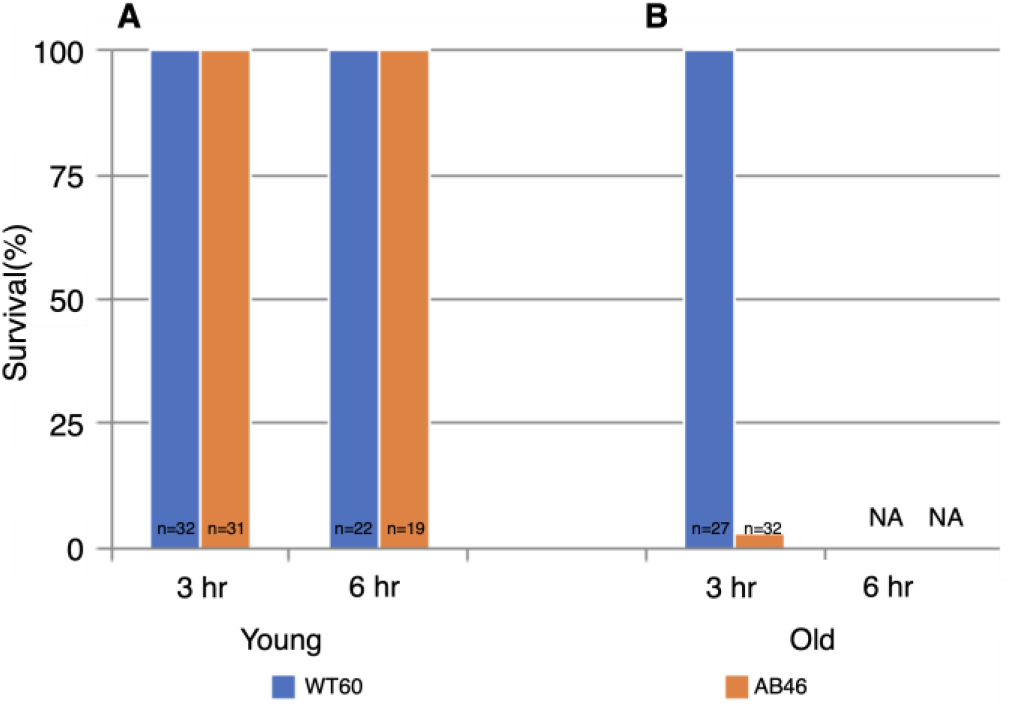
Survival Following Prolonged Hypoxia. Young flies (panel A) or old flies (panel B) were subjected to either 3 hours or 6 hours of anoxia using a 25mL/sec flow rate of nitrogen. Flies were then returned to standard atmosphere and monitored for recovery from the spreading depression coma as described in the legend of Figure 2. Animals that did not recover movement within 5 hours were scored as not surviving the hypoxia. NA indicates that none of the animals survived.

## 4. Discussion

Although the role of MSR in oxidative stress has been extensively studied, the investigation into how the absence of MSR activity affects hypoxia tolerance in *Drosophila melanogaster* is a relatively new area of exploration. Fruit flies were discovered to be tolerant to acute hypoxia (0mm Hg O_2_) in the early 1990s, where they survived in oxygen depleted environments for several hours without any evidence of injury [1]. The disruption of oxygen homeostasis is a major factor for many disease etiologies and pathobiology [17]. The low oxygen conditions (hypoxia) used in this study is a prominent clinical problem associated with many diseases such as ischemic heart disease, cerebral ischemia (stroke), pulmonary hypertension, diabetes complication, high altitude illness, and cardio-respiratory disorders (bronchopulmonary dysplasia and obstructive sleep apnea). The evolutionary conservation of genetic and signaling pathways from *Drosophila* to mammals allows for it to be an ideal model system to investigate the genetic basis of hypoxia tolerance [17]. Fruit flies enter a protective coma called spreading depression, which allows survival to prolonged periods of hypoxia, by suppressing their overall metabolic rate to prevent cellular injury during reoxygenation [18]. Hypoxia/reoxygenation induces cellular injury through the promotion of oxidative stress, which was the impetus to quantify recovery during these conditions. ROS cause oxidative damage to amino acids, lipids, nucleic acids, and play a crucial role in aging and senescence [19]. The MSR system protects vital cell constituents from free radical damage and counteracts oxidative stress [20].

The loss of all known MSR activity has been reported in bacteria [21] and yeast [22], although these organisms are not as developmentally complex as *Drosophila*. The correlation between the loss of MSRA activity and reduction in longevity were documented in yeast [22], *C. elegans* [23] and mice [24]. However, a recent study on an *MSRA* knockout mouse did not find an effect on lifespan [25]. In our lab, *MSRA* and *MSRB* gene deletions were created through imprecise excision of p-element transposons located in each gene, leading to the establishment of *Drosophila* as the first known *in-vivo* animal model to lack any known MSR activity. We have previously found a significantly shorter lifespan in the *MSR* double mutant with nearly full restoration of normal lifespan in the presence of a single wild-type allele of either *MSRA* or *MSRB* [13]. Our previous experiments failed to identify a significant phenotype in *Drosophila* lacking just one of the two *MSR* genes. In fact, there is a significantly longer third instar with larvae growing at a slower rate and adult flies having a shortened lifespan in the absence of any MSR activity (*MSRA*^Δ/Δ^*MSRB*^Δ/Δ^) [13]. Similarly, we found key behavioral differences on the effects of hypoxia under conditions that vary the rate which the spreading depression coma is induced as well as the length of hypoxia stress.

The time required for flies to cease movement due to the protective coma induced by hypoxia can be varied by altering the rate at which the chamber becomes hypoxic (i.e., by controlling the flow rate of the nitrogen gas). After sudden exposure to hypoxia, the *MSR* double mutant flies took significantly longer to recover compared to the wildtype flies throughout the flies’ entire lifespan (Figure 1). More interestingly, the MSR-deficient flies showed a markedly reduced ability to survive the hypoxic stress as they approached senescence with only 38% surviving among the flies that were 40-45 days old (Figure 4). In contrast, the wildtype strains had nearly 100% survival of the hypoxia throughout their entire lifespan including the period of senescence (60-65 days old) (data not shown). According to the oxidative stress theory of aging, as animals age, an increase in ROS and oxidative stress plays a role in governing lifespan. Previous research suggests that the accumulation of oxidative damage as part of the aging process, in addition to ROS produced during hypoxic stress, leads to increased cellular damage, which results in a longer recovery time and decreased percentage of survival as the flies age [26,27]. This may also be due to the reduction in the levels of both isoforms of MSR enzymatic activity with aging [28]. These experiments are the first evidence of an age-dependent effect of MSR deficiency in both recovery and survival from hypoxic stress.

Being that we measured hypoxia tolerance through percent survival and/or recovery time, we were curious to see whether this would be altered if spreading depression was induced more slowly. The onset of spreading depression was not affected by the lack of MSR activity but there was an age-dependent decrease in the time to induce the coma for both the moderate (Figure 2A) and slow (Figure 2B) flow rates. However, upon reoxygenation, the MSR-deficient flies displayed a stark contrast to previous experiments and recovered faster than wildtype flies throughout their entire lifespan (Figure 3). More interestingly, the survival of the MSR-deficient flies improved dramatically and was nearly 100% throughout the entire lifespan (Figure 4). Our results suggest slow induction of spreading depression may allow for ischemic preconditioning to commence. Mild ischemic stress triggers late preconditioning through the release of chemical signals, referred to as “triggers.” These triggers in the form of nitric oxide, reactive oxygen species, and adenosine activate a signal transduction cascade through protein kinase C (PKC) and Janus-activated kinases 1 and 2 (JAK1/2), leading to the activation of cytoplasmic as well as stress-responsive transcription factors including NF-kB, STAT1, and STAT3. Downstream events result in the upregulation of cardioprotective genes containing the inducible isoform of NOS (iNOS), cyclooxygenase (COX)-2, and antioxidant enzymes such as SOD. Thus, neuroprotective signaling and decreased apoptotic pathway activation as well as gene reprograming and metabolic downregulation may be why MSR-deficient flies recover faster and have markedly improved survival when the induction of hypoxia occurs more slowly [29].

The faster recovery and improved survival following hypoxia in the MSR-deficient flies upon slow induction of spreading depression may be a result of perturbations in the serotonergic system. The serotonergic neurons in *D. melanogaster* are responsible for the coordination of feeding behaviors and body mass control through the insulin/IGF pathway (IIS) [30]. Nuclear translocation of transcription factor daf-16 following the genetic ablation of tryptophan-hydroxylase, a critical enzyme involved in serotonin biosynthesis, has been reported in *C.elegans [31].* The *Drosophila* homolog for daf-16 is dFOXO and is negatively controlled through Ser/Thr phosphorylation via the induction of the insulin/IGF signaling pathway (IlS) [32]. It has been demonstrated that dFOXO activity has a direct effect on MSR transcriptional activation [33]. *dFOXO* deletion mutants express multiple phenotypes that are consistent with MSR deficient flies including decreased survivability, delayed larval development, and reduced fecundity [34]. *dFOXO* deletion mutants also show a decreased post hypoxic survivability, as we are seeing in senescent *MSR* deletion mutants under rapid hypoxic induction. The effect of hypoxic induction rate has not been reported in the context of *dFOXO* loss of function mutations. It has been implicated that dFOXO and its involvement in hypoxic protection involves known immune transcription factor NF-kB/*relish* [35]. MSR may also then be involved in a concerted mechanism with dFOXO and *relish* or in a disparate, yet related pathway. Altered serotonergic signaling can inhibit the insulin/IGF pathway, resulting in developmental delay and growth retardation [36]. We propose the probability that mutations in the insulin/IGF pathway of *Drosophila* display deleterious effects on longevity, resistance to oxidative stress, and muscle development due to alterations in the mTORC1 pathway as well as potentially dysregulated *MSR* transcription [37].

The hypoxia tolerance in fruit flies permits survival of extended hypoxia without neuronal deficit, due to the protective coma they entered during hypoxia [38]. Our next set of experiments focused on studying hypoxic tolerance upon prolonged hypoxia (3 and 6 hours) and the slow induction of spreading depression. Young animals had 100% survival for both the wild type and MSR-deficient strains up to six hours of slow induction of hypoxia (Figure 6). At old age, 100% of the wildtype flies survived three hours of hypoxia although none of the animals survived six hours of hypoxia (Figure 6B). In sharp contrast, only 3% (1 of 32 flies) of the old MSR-deficient flies survived the three hours of hypoxia (Figure 6B). There is a clear age-dependent decline in the ability to survive prolonged hypoxia in the absence of MSR. Young MSR-deficient flies continued to recover significantly faster than wildtype animals upon prolonged hypoxia, while old MSR-deficient flies did not recover at all (Figure 5). The underlying mechanism may be associated with the relationship between the production and depletion of cellular energy during spreading depression. During hypoxia, the metabolic rate is known to significantly decrease to allow *Drosophila* to preserve cellular ATP while also decreasing its total production [15,39–41]. When flies start to recover upon reoxygenation, there is less available ATP to restore metabolic deficits due to the ATP depletion during the period of hypoxia. Overall, survival is compromised [39,42]. In addition, ATP depletion is known to lead to failure of the Na^+^/K^+^ ATPase, which results in dysregulation of ionic homeostasis, protein unfolding and subsequently protein aggregation [43]. Numerous reports in the literature reflect similar trends of a strong inverse correlation between increased stress duration and decreased survival probability, possibly due to deficiency in ATP production and the inability of the fly to compensate for ATP consumption [15,41].

## 5. Conclusions

The results obtained from this study demonstrate that MSRA and MSRB play an age-dependent role in protection against oxidative stress throughout the lifespan of *D. melanogaster*. MSRA and MSRB are known to behave as antioxidants to reduce methionine sulfoxide (nonfunctional form of methionine from ROS oxidation) back to the functional form of methionine [44]. The original expectation was that MSR-deficient flies would have a compromised ability to tolerate hypoxia. The results of experiments using the MSR-deficient flies suggest new lines of inquiry involving ischemic preconditioning, longevity, and serotonin pathways. These results support previous studies that suggest the activation of protective mechanisms to defend against oxidative stress, essentially leading us to a better understanding how these *MSR* genes affect aging. Our studies offer possible insight into hypoxic-like conditions in humans, such as stroke, that may ultimately contribute to better drug design or other treatments.

## Author Contributions

Conceptualization, Nirthieca Suthakaran and David Binninger; Data curation, Sanjana Chandran and Michael Iacobelli; Formal analysis, Nirthieca Suthakaran and David Binninger; Funding acquisition, David Binninger; Investigation, Nirthieca Suthakaran and David Binninger; Methodology, Nirthieca Suthakaran and David Binninger; Project administration, David Binninger; Resources, David Binninger; Supervision, David Binninger; Writing – original draft, Nirthieca Suthakaran, Sanjana Chandran and Michael Iacobelli; Writing – review & editing, Nirthieca Suthakaran, Sanjana Chandran, Michael Iacobelli and David Binninger.

## Acknowledgments

I (D.B.) would like to thank Ken Dawson-Scully at Florida Atlantic University (FAU, Boca Raton, FL) for suggesting that I investigate whether MSR affects hypoxia tolerance in *Drosophila*. I would like to thank the numerous undergraduate students who have provided laboratory support to maintain the fly stocks and assist with some of the experiments described in this paper. I would like to also thank Kelsey Binninger for her assistance with the graphics. Finally, I am grateful to my longtime colleague and friend Herbert Weissbach who introduced me to the MSR system and has been continuously supportive of my studies.

